# Soil fungal communities vary more with soil characteristics than tree diversity at a local scale

**DOI:** 10.1101/2022.01.26.477882

**Authors:** Steve Kutos, Elle M. Barnes, J.D. Lewis

## Abstract

Soil fungal communities vary spatially due to factors including variations in plant diversity and soil characteristics; however, the relative influences of these factors on composition and therefore function remain unclear. Small-scale variation in fungal communities may drive local variation in nutrient cycling and decomposition and may respond more to local factors compared with large climatic variations. Clarifying the roles of these factors can improve our predictions of soil fungal community and biogeochemical cycling responses to anthropogenic changes. Therefore, we examined relationships among abiotic and biotic factors and soil fungal communities associated with *Quercus rubra* and *Pinus resinosa* saplings and mature trees in a mixed-hardwood woodland. We also compared community composition and fungal enzymatic activity. Fungal community composition was associated with spatial heterogeneity of soil characteristics, while host sapling and tree species identity were poor predictors of community composition. Further, most of the compositional variation was unexplained by measured variables, suggesting stochasticity and other environmental characteristics may drive spatial variation in these communities. Additionally, enzymatic activity did not clearly correlate with fungal community composition. Overall, soil fungal communities and enzymatic activity adjacent to saplings in this woodland are influenced primarily by soil characteristics and stochasticity and not plant identity.

## INTRODUCTION

Soil fungal communities provide essential ecosystem services, yet we are still learning how and why these communities are distributed across space (Bardgett and van der Putten 2014; Tedersoo et al. 2014; Guerra et al. 2020). Robust spatial mapping of soil fungal taxa has the potential to inform us of their functional roles, environmental and perturbation tolerances, and allow us to make stronger predictions of how these taxa may be impacted by future environmental change (Ettema and Wardle 2002). Broad evidence suggests that soil fungal biogeography is not random but displays distinct spatial patterns (Talbot et al. 2014; Tedersoo et al. 2014; Lladó et al. 2018). However, disparities remain in our understanding of the driving factors of these fungal biogeographic patterns (Bahram et al. 2016; Tedersoo et al. 2014; Hendershot et al. 2017; van der Linde et al. 2018; Guerra et al. 2020).

In temperate forests, soil fungal communities are influenced by many biotic and abiotic factors (Talbot et al. 2014; Tedersoo et al. 2014; Lladó et al. 2018; van der Linde et al. 2018). Biotic factors can include the forest successional stage, leaf litter quality, and aboveground plant distributions (Talbot et al. 2014; Baldrian 2017; Lladó et al. 2018). This last factor has warranted consideration because of the increasing evidence of the role of plant species identity and diversity on the distribution of soil fungal taxa (Ishida et al. 2007; Tedersoo et al. 2008; Urbanová et al. 2015; Leff et al. 2018). This evidence suggests that at least some soil fungal taxa can exhibit a preferential association for a narrow range of plant species (Dickie 2007; Urbanová et al. 2015; Leff et al. 2018). For example, forest trees can influence fungal community composition in the rhizosphere and local bulk soil by their canopy cover density, the release of root exudates, and their litter-fall, which can all create gradients of soil chemistry (Clarholm and Skyllberg 2013; Prescott and Grayston 2013). In these forests, mature trees can also function as a reservoir of fungal inoculum for closely developing seedlings (Cline et al. 2005; Dickie and Reich 2005; Lang et al. 2011). Therefore, the establishment and growth of tree seedlings as well as its soil fungal community could depend on the microsite at which they establish (Tedersoo et al. 2008). The biotic factors can also interact directly and indirectly with soil abiotic characteristics such as pH, nutrient concentrations, moisture, and heavy metal inputs, which can lead to further dissimilarity in fungal community composition (Tedersoo et al. 2014; Essene et al. 2017; Lladó et al. 2018; Van Geel et al. 2018). Despite decades of research, a knowledge gap remains in how each of these abiotic and biotic factors affects soil fungal community composition (Lladó et al. 2018; Guerra et al. 2020). This gap is further confounded by research scale, as the factors that influence soil fungal composition on continental scales can be different than on regional or smaller local scales (Bahram et al. 2016; Lladó et al. 2018; Guerra et al. 2020).

On local scales, such as in small forests or woodlands, large spatial heterogeneity of abiotic and biotic factors can decline, possibly leading to increased fungal community similarity (Ettema and Wardle 2002; Lekberg et al. 2007; Peay and Bruns 2014; Baldrian 2017). Since broad geographic barriers are likely decreased at the local scale, the importance of fungal spore dispersal may become increased as the success of fungal spore colonization can depend on both soil characteristics and local plant-host diversity (Lekberg et al. 2007; Peay et al. 2012; Peay and Bruns 2014). Understanding the relative influence of these factors might tell us how fungal taxa are distributed through space more broadly as well as the roles of environmental filters in local woodlands (Lladó et al. 2018; Guerra et al. 2020).

To explore these themes, we examined the relative influence of tree species identity, tree compositions, and spatial heterogeneity of soil characteristics on soil fungal communities of tree saplings in a small woodland in Armonk, NY, USA. A field experimental assay was employed using saplings of two common eastern US trees (*Pinus resinosa* and *Quercus rubra*) planted alone and planted together around two mature tree species (*Quercus rubra* and *Fagus grandifolia*) distributed throughout this woodland. We hypothesized that differences in fungal community composition would be driven by both spatial heterogeneity of soil characteristics and nearby mature tree species identity regardless of sapling species or composition, as they can obtain their mainly later-successional fungal taxa from those associating with nearby trees (Tedersoo et al. 2008). Additionally, we assessed if potential changes in fungal community composition correlates with differences in the activity of extracellular enzymes associated with local carbon and phosphorus cycling. For this, we hypothesized that soil enzymatic activity will not differ between the treatments but will differ between the two different mature tree species and woodland locations.

## MATERIALS AND METHODS

### Study site and experimental design

To explore our hypotheses, we utilized a field experiment in the woodland at Fordham University’s Louis Calder Center in Armonk, NY, USA (41.131789, -73.732911). The tree diversity of this 110-acre woodland is mainly composed of oaks (*Quercus* spp.), maples (*Acer* spp.), American beech (*Fagus grandifolia)*, and hickory (*Carya* spp.). Despite being in a suburban setting near New York City, this woodland has been relatively unmanaged for over a century. However, like many woodlands in the region, it has a sizeable quantity of deer browse which reduces the low vegetation layers. Climate is temperate with mean annual temperatures of ∼12°C and mean annual precipitation of ∼120 cm. Bedrock parent material consists of a combination of granite, gneiss, and marble, with soils designated as acidic sandy loams (Schuberth 1968; Edinger 2014). To utilize differences in soil characteristics across this woodland, four area blocks of comparable size were established. Within each of these four blocks, mature trees of *Quercus rubra* and *Fagus grandifolia* were randomly selected for use within the field assay (a total of 10 trees per species throughout all blocks). These four blocks were numbered in the order of a slight elevation decline: *Block 1* is an area at the top of a lakeside ridge, *Block 2* and *Block 3* are ∼40-50 m down the small ridge from *Block 1* with increased soil moisture due to precipitation runoff from *Block 1*, and *Block 4* is a flat area at the bottom of the ridge with a nearby vernal pool. Total area of the woodland assay was ∼75 acres.

One-year-old bareroot saplings of northern red oak (*Q. rubra*) and red pine (*Pinus resinosa*) were obtained from Cold Stream Farm (Freesoil, MI, USA) and immediately planted in sterilized 3.8 cm cone-tainer pots (Stuewe and Sons Inc., Tanget, OR, USA) with field-collected soils from underneath corresponding trees of the same species found within the Calder Center woodland. These sapling species were chosen because they are common in eastern US forests, fast-growing, and are associated with ectomycorrhizal fungal taxa. In early spring 2019, from this stock, saplings were randomly selected to create three treatments which were planted around each of the 20 mature trees within the woodland: (1) a solo-planted *Q. rubra* sapling, (2) a solo-planted *P. resinosa* sapling, and (3) a paired planting of *P. resinosa* and *Q. rubra* saplings ∼30 cm apart. For these treatments, each sapling was planted ∼1.5 m from the bole of the mature tree at randomly chosen cardinal directions. The cardinal direction without a planted sapling was designated as a no-sapling control treatment. Overall, there were 60 total sapling planted treatments and 20 control treatments throughout the woodland. Planted tree saplings were protected from deer browse using wire fences giving at least a 1-foot barrier between fence and sapling to not obstruct growth. Soil moisture and pH were evaluated monthly around each treatment with pH measured using a 2:1 ratio of distilled water to soil on an Accumet AE150 probe (ThermoFisher Scientific, Waltham, MA, USA) and soil moisture measured by drying 5 g of field moist soil for 24 h at 105°C.

### Soil sampling, DNA sequencing, bioinformatics, and statistical analysis

In late autumn 2019, soil cores were collected ∼10 cm in depth at three points in close proximity around each of the planted and control treatments. These soils were merged and sieved through a sterilized screen and stored at -20° C. Fungal DNA was extracted from 0.25 g of soil using the PowerSoil DNA Isolation Kit (Qiagen Inc., Mississauga, Canada). PCR amplification targeted the forward primer ITS1F and reverse primer ITS2 (Smith and Peay 2014) with 25 μl reactions containing: 2.5 μl 10X Invitrogen buffer, 0.75 μl 50 mM MgCl_2_, 1.0 μl 10 mM of both primers, 0.5 μl 10 mM dNTPs, 0.25 μl Platinum Taq (5 U/μl), 1.0 μl BSA, and 2 μl DNA. These amplifications were conducted on an Applied Biosystems thermocycler (Model 2720, Foster City, CA, USA) under the following conditions: 94° C for 1 min, followed by 94° C for 30 sec, 58° C for 30 sec, 68° C for 30 sec for 35 cycles, and a final extension at 68° C for 7 min. These amplicons were purified using Sera-Mag Speedbeads™ (GE Healthcare, Chicago, IL, USA) and quantified with a Qubit 4.0 Fluorometer (ThermoFisher, Invitrogen, Carlsbad, CA, USA). These libraries, including negative and positive controls, were normalized, pooled, and sequenced on a 2 × 250 bp Illumina MiSeq at Genewiz (Brooks Life Sciences Company, South Plainfield, NJ, USA). The positive sequencing control contained ten known fungal taxa from the phyla Basidiomycota, Ascomycota, and Mortierellomycota. Demultiplexed FASTQ files were filtered and processed using a QIIME2 bioinformatic pipeline (Release: 2020.8; Caporaso et al. 2010). Sequencing adapters, primers, and low-quality bases were trimmed using *ITSXpress* and *cutadapt* (Martin 2011; Rivers et al. 2018). Sequences were aligned and merged through DADA2 to produce ASVs (Callahan et al. 2016) and assigned to taxonomic groups using the UNITE database (Version 8.2; Abarenkov et al. 2010). Samples were rarified to a read count depth of 1335. Finally, FUNGuild was run on these taxonomic assignments to attach functional groupings using only probable and highly probable assignments (Nguyen et al. 2016).

All statistical synthesis was completed in R (Version 4.0.1; Oksanen et al. 2007). Alpha diversity was measured using the Shannon diversity index and ASV richness, with mixed-effect linear models used to evaluate for significant differences. Beta diversity was measured on log-transformed read counts using Bray-Curtis dissimilarity. Homogeneity of dispersions was checked using *betadisper* and PERMANOVAs with a blocking effect were run to test for significance between the treatments and the woodland locations with an NMDS ordination used for visualization. Indicator species analysis was performed with a correction for unequal sample size to examine which ASV significantly associated with each treatment using the *indicspecies* package with 9999 Monte Carlo permutations (De Caceres et al. 2016).

### Enzymatic fluorometric assay and soil elemental analysis

Within three days of soil collection, enzymatic potential activity of six enzymes common in soils within the samples was evaluated using a high throughput fluorometric assay (Bell et al. 2013; supplementary material S1). From each sample, a slurry was created by mixing 2.75 g of soil and 91 ml of 50 mM sodium acetate buffer. These slurries were incubated for three hours in the dark at room temperature (∼23° C) in 96 deep-well plates before being centrifuged for 30 mins at 2300 x g. Supernatants of 200 ul were transferred to black flat-bottom plates and measured on a SpectraMax M2e microplate reader (Molecular Devices, San Jose, CA, USA) with the excitation wavelength at 365 nm and the emission wavelength at 450 nm. A small amount (10 ul) of NaOH was added prior to plate measurement to increase activity of the substrates. A control plate consisting of only buffer and substrate was used for each sample. These controls were applied to the raw values, and then activities were converted to nmol h^−1^ g^−1^. Total potential enzymatic activity was compared among treatments on log-transformed values using a mixed-effects linear model.

Soil analyses were completed to evaluate relationships between soil characteristics and soil fungal community composition. Soil samples were analyzed with inductively coupled mass spectrometry of 13 elements, including total percent of nitrogen and carbon, at the Cornell Nutrient Analysis Laboratory (Ithaca, NY, USA). Elemental raw values were log-transformed and compared using a mixed-effect linear model. Spatial relationships between fungal community composition and these soil characteristics were assessed using redundancy analysis (RDA). Matrices were log-transformed prior to running the RDA to not overweight the considerable number of zeros in the data, and model significance was determined using an ANOVA. Finally, to examine the relationship between fungal community dissimilarity, soil characteristic differences, and spatial distance, a partial Mantel test was run using Pearson correlations on the full community and the most common fungal phyla. For this partial Mantel, the soil characteristic matrix and ASV matrix were log-transformed, and Bray-Curtis dissimilarity distance was used.

## RESULTS

### Soil fungal community composition patterns

After all filtering steps, data analysis identified 1,686 fungal ASVs from 41 samples (remaining samples were removed from analysis due to low read counts or mortality during experimental assay period). The fungal community was predominantly composed of taxa from the phylum Basidiomycota (∼66 % of all reads); lesser numbers of ASVs were observed from Ascomycota (∼17 %) and Mortierellomycota (∼9 %, Fig. 1). The ten most abundant ASVs in all treatments were taxa from the phyla Basidiomycota (Order: Tremellales, Russulales, Cantharellales, Agaricales, Atheliales) and Mortierellomycota (Order: Mortierellales). There were no ASVs from any phyla that were found in all samples. Soil fungal communities among the planted sapling treatments (solo pine, solo oak, or paired pine and oak) and no-sapling control treatments did not differ significantly in either Shannon diversity (*F* = 0.11, *p* = 0.94, Fig. 2) or ASV richness (*F* = 0.35, *p* = 0.8). Further, there were no significant differences in fungal community composition among the planted saplings and no-sapling control treatments using Bray-Curtis distance (*F* = 0.91, *p* = 0.83). The soil fungal communities between all treatments around either mature tree *Quercus rubra* or *Fagus grandifolia* were also not significantly different in Shannon diversity (*F* = 0.57, *p* = 0.46) or ASV richness (*F* = 2.03, *p* = 0.17). However, these communities did significantly differ in beta diversity using Bray-Curtis distances (*F* = 1.8, *p* = 0.004, r^2^ = 0.04). However, it is important to note the low *r*^*2*^ value. As expected, the fungal communities were significantly different among different areas of the woodland in Shannon diversity (*F* = 5.6, *p* = 0.012; Fig. 2) and ASV richness (*F* = 6.85, *p* = 0.006). The composition of these communities was also significantly different in Bray-Curtis distance (*F* = 2.14, *p* < 0.001, r^2^ = 0.12; Fig. 2).

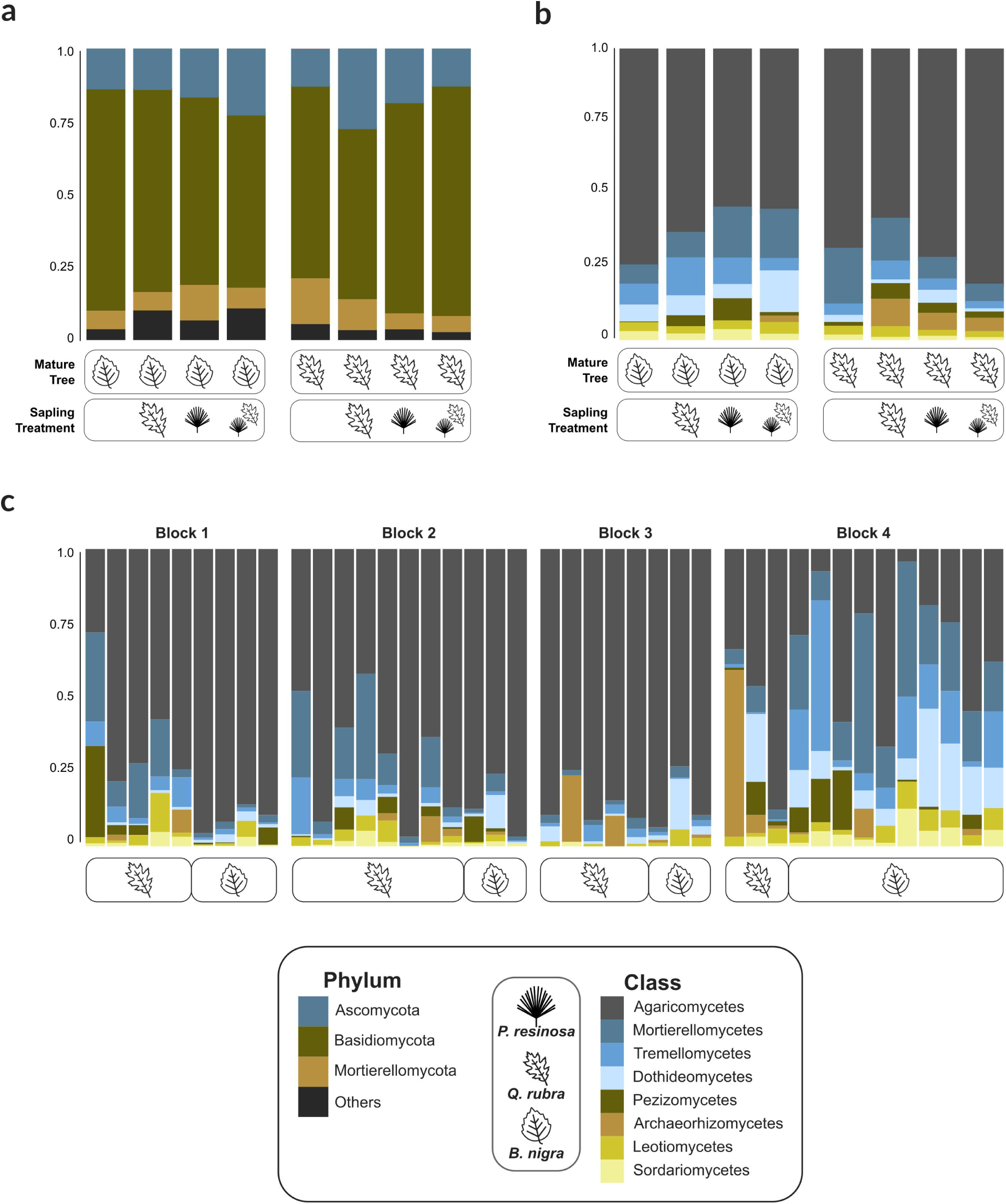

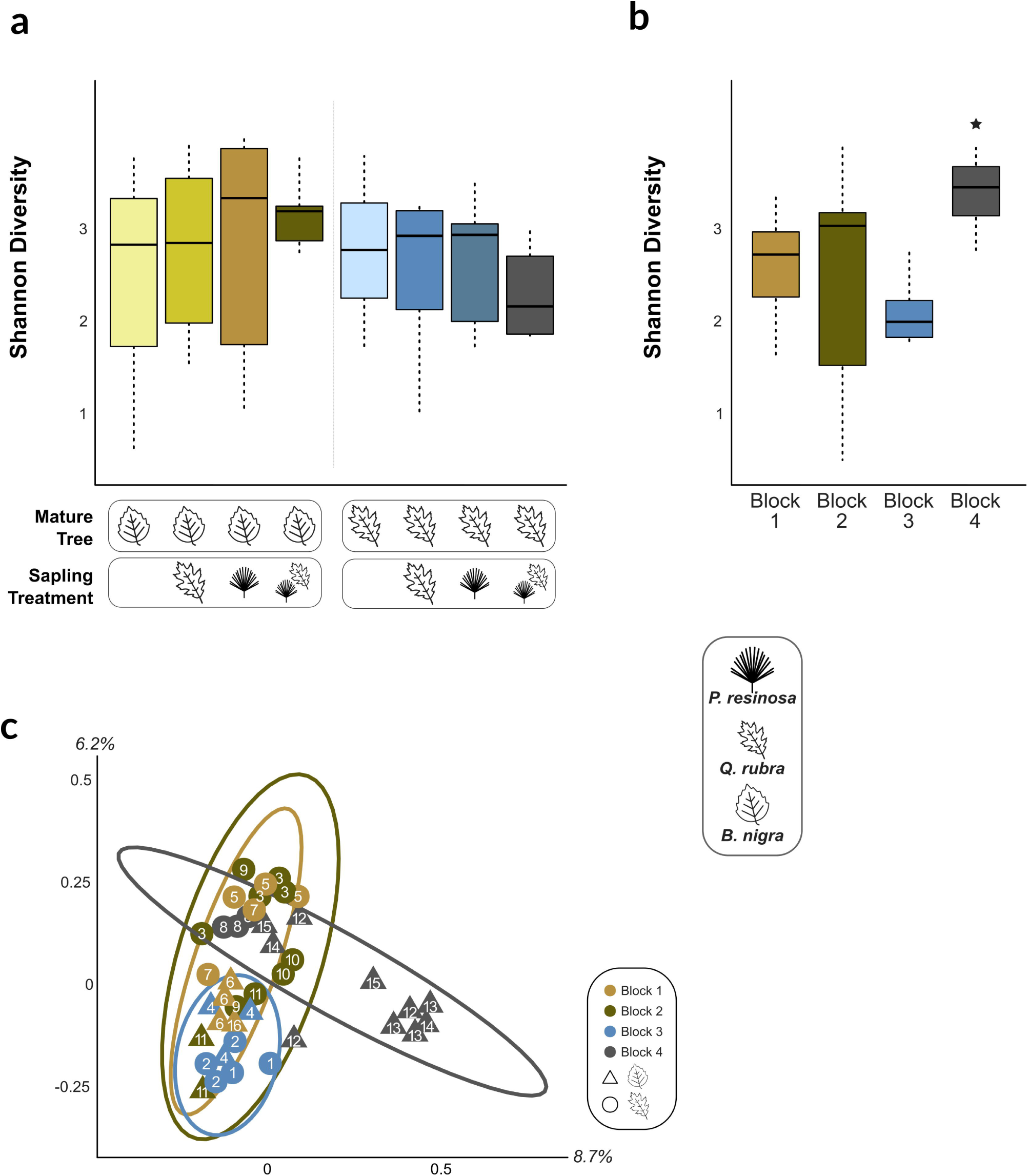

Taxa classified in FUNGuild as ectomycorrhizal taxa were not significantly different in Shannon diversity or ASV Richness between the planted sapling treatments, treatments around either *Quercus rubra* or *Fagus grandifolia*, or different areas of the woodland (*p* > 0.05) but were significantly different in Bray-Curtis distance between the different areas of the woodland (*F* = 1.4, *p* = 0.002, r^2^ = 0.12). Taxa classified in FUNGuild as saprobic taxa were also not significantly different in Shannon diversity or ASV Richness between the planted sapling treatments, treatments around either *Quercus rubra* or *Fagus grandifolia*, or different areas of the woodland (*p* > 0.05). These taxa were significantly different in Bray-Curtis distance between the different areas of the woodland (*F* = 1.8, *p* < 0.001, r^2^ = 0.13). It is important to note that the relative abundance of these taxa classified as either ectomycorrhizal or saprobic via FUNGuild were low.

### Fungal taxa relative abundance patterns

Within the dataset at the phyla level there was not a clear difference in broad relative abundance patterns between planted sapling and no-sapling control treatments (Fig. 1). This pattern matches the lack of significant difference in alpha and beta diversity among these treatments. Overall, all treatments had greater relative abundances of the fungal family Russulaceae (Phylum: Basidiomycota; Class: Agaricomycetes) than all other families. Of note, taxa in this family were ∼3X lower in relative abundance in the no-planted sapling treatments compared to all other treatments. Thus, their relative abundance was higher if a sapling was nearby. Other planted sapling treatments had distinct patterns in other abundant fungal families. For instance, taxa in the fungal family Sebacinaceae (Phylum: Basidiomycota; Class: Agaricomycetes) were ∼10X higher in relative abundance when near a mature *Quercus rubra*. Oppositely, taxa in the fungal family Mycosphaerellaceae (Phylum: Ascomycota; Class: Dothideomycetes) were ∼8.5X higher in relative abundance when near a mature *Fagus grandifolia*. These relative fungal relative abundance patterns of the mature trees were also found among the different sapling species. For example, *P. resinosa* saplings near *F. grandifolia* had a ∼15X higher relative abundance of taxa in the fungal family Hydnaceae than when planted around *Q. rubra*.

There were also distinct relative abundance and diversity patterns spatially throughout the woodland (Fig. 1C). The dryer area on the top of the ridge (*Block 1*) had high relative abundances of the fungal families Russulaceae (∼49 % of reads), Sebacinaceae (∼11 %), and Mortierellaceae (∼9 %). The highest relative abundant family of the area down the ridge (*Block 2)* was also Russulaceae (∼23 % of reads), along with the families Hydnaceae (∼15 %), and Mortierellaceae (∼10 %). The area down the ridge and to the south of Block 2 (*Block 3)* also had a high relative abundance of Russulaceae (∼56 % of reads) with lower abundances of all other families. Finally, the area at the bottom of the ridge which receives heavy precipitation runoff (*Block 4*) slightly diverges from these diversity patterns with high relative abundances in the families Russulaceae (∼14 % of reads), Mortierellaceae (∼13 %), and Trimorphomycetaceae (∼6 %). Therefore, in this area, taxa in the family Russulaceae were 2X – 5X lower in relative abundance than the other woodland area blocks. Additionally, taxa in the family Piskurozymaceae only had high relative abundances in this area when near a *F. grandifolia*.

Indicator species analysis was used to identify ASVs that may significantly associate with the planted sapling treatments and between treatments around *Q. rubra* or *F. grandifolia*. Throughout all ASVs in the dataset, none were significant as indicator taxa among the planted saplings (solo pine, solo oak, or paired pine and oak). There were three ASVs there were significant (*p* < 0.01) as indicator taxa around *F. grandifolia* including two ASVs in families in the phylum Ascomycota (Mycosphaerellaceae and Cucurbitariaceae) and an ASV in the family Ganodermataceae (Phylum: Basidiomycota; Class: Agaricomycetes). Similarly, three ASVs there were significant (*p* < 0.01) as indicator taxa around *Q. rubra* including two ASVs the family Mortierellaceae (Phylum: Mortierellomycota) and a unideitifed ASV in the class Saccharomycetes (Phylum: Ascomycota). Of note, the relative abundance of these ASVs were low in the respective treatments.

### Enzymatic activity and soil characteristics

There was no significant difference between any of the planted sapling treatments or woodland locations in activity of five enzymes: acid phosphatase, cellobiohydrolase, β-glucosidase, α-glucosidase, and β-xylosidase (supplementary material S2). Additionally, there was no discernible activity of β-glucuronidase and therefore was removed from analysis. While not significantly different, there were on average higher enzymatic activities in treatments around *Q. rubra* than *F. grandifolia* (β-glucosidase: ∼1.5X higher, acid phosphatase: ∼1.3X higher, β-xylosidase: ∼1.2X higher, and cellobiohydrolase: ∼1.8X higher; supplementary material S2) as well as on average lower activities in the area of the woodland at the bottom of the ridge (*Block 4*) compared to the other woodland areas (β-glucosidase: ∼1.4X lower, acid phosphatase: ∼1.5X lower, and α-glucosidase: ∼1.5X lower; supplementary material S2).

Of the soil characteristics examined, there were no significant differences between the different planted sapling treatments or between the soils around *Q. rubra* or *F. grandifolia*. Concentrations of several elements differed spatially and significantly among locations within the woodland including soil moisture (*F* = 4.3, *p* = 0.03), Zn (*F* = 7.6, *p* = 0.004), and Na (*F* = 5.4, *p* = 0.01). This pattern was further shown in the RDA where predictive soil characteristics were significantly correlated with fungal community composition (*F* = 1.2, *p* < 0.001; Fig. 3). Overall, these characteristics explained ∼33% of the total proportion of constrained variation in the model. However, the adjusted R^2^ only explained ∼8 % of the total variation in the data.

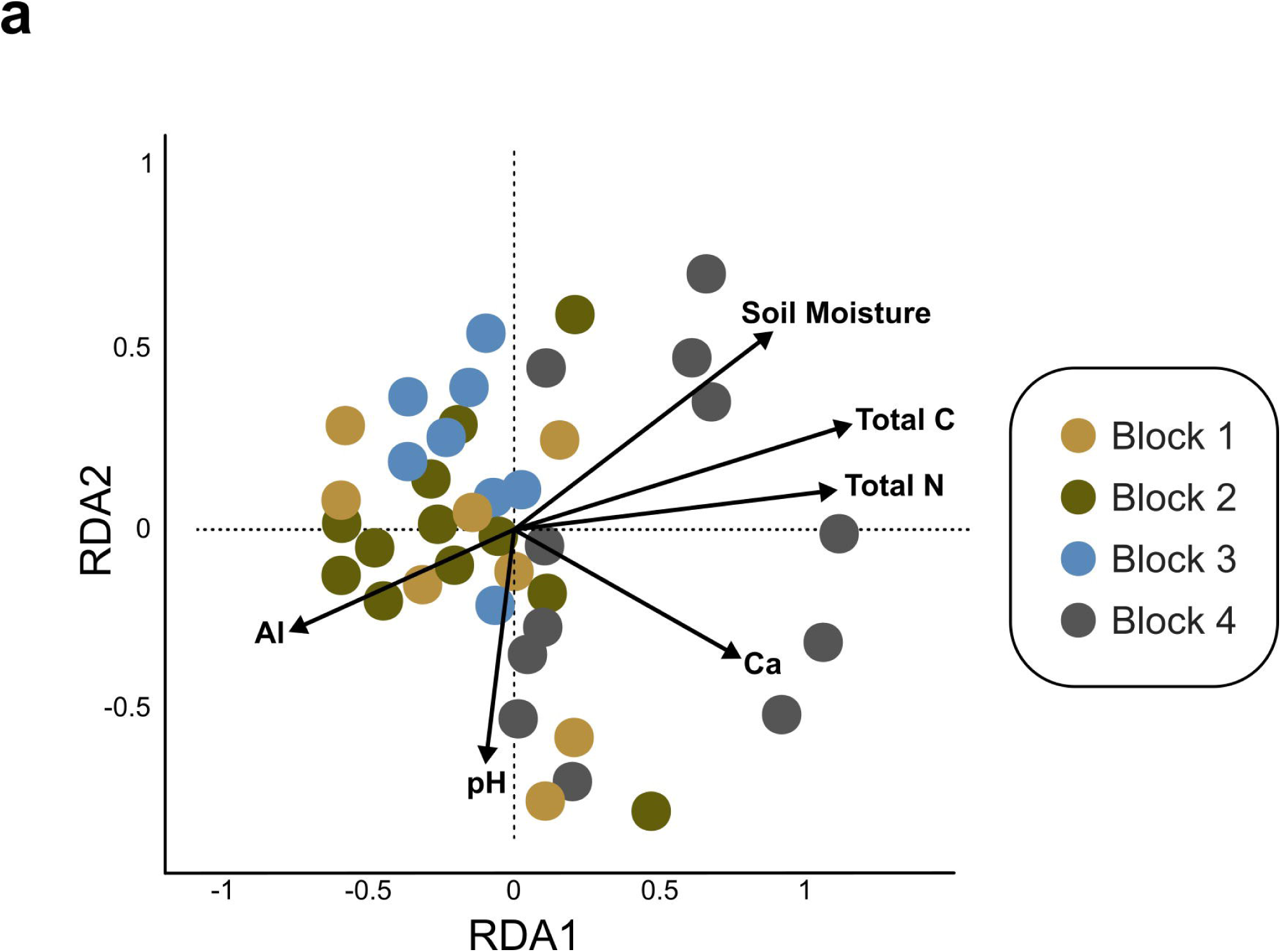

Fungal ASVs were compared with these soil elements and characteristics via a partial Mantel test controlling for distance within the woodland (Table 1). Soil fungal community composition was significantly associated with differences in total N % (r = 0.26, *p* = 0.005), soil moisture (r = 0.26, *p* = 0.003), Al (r = 0.33, *p* = 0.006), total C % (r = 0.19, *p* = 0.017), and pH (r = 0.12, *p* = 0.02). Distinct soil characteristic association patterns were found when focusing on the three dominant phyla found in the data and the two functional groups (Table 1). Basidiomycota taxa were only significantly associated the with Al (r = 0.22, *p* = 0.014). Similarly, Ascomycota taxa were only significantly associated the Al (r = 0.16, *p* = 0.03). Mortierellomycota taxa were significantly associated with Al, soil moisture, total C % and N %, and pH (Table 1).

**Table 1.**
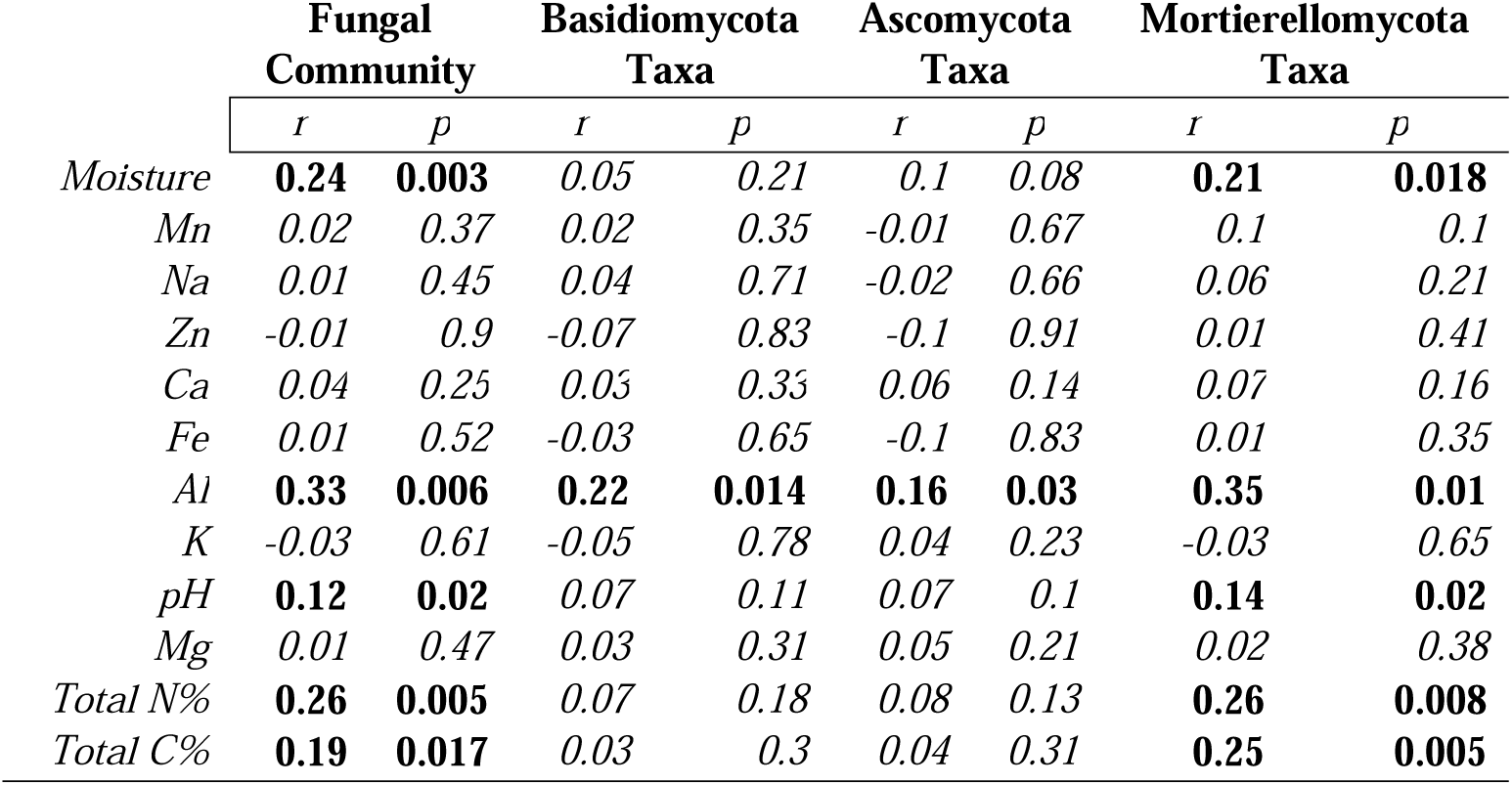
Partial Mantel. Pearson correlations using Bray-Curtis distances between soil fungal composition of the fungal community and high-relative abundant phyla and physicochemical variables. ASV and elemental matrices were log-transformed. Bold denotes significant Pearson correlation at *p* < 0.05.

|  | Fungal Community |  | Basidiomycota Taxa |  | Ascomycota Taxa |  | Mortierellomycota Taxa |  |
| --- | --- | --- | --- | --- | --- | --- | --- | --- |
|  | *r* | *p* | *r* | *p* | *r* | *p* | *r* | *p* |
| *Moisture* | **0.24** | **0.003** | 0.05 | 0.21 | 0.1 | 0.08 | **0.21** | **0.018** |
| *Mn* | 0.02 | 0.37 | 0.02 | 0.35 | -0.01 | 0.67 | 0.1 | 0.1 |
| *Na* | 0.01 | 0.45 | 0.04 | 0.71 | -0.02 | 0.66 | 0.06 | 0.21 |
| *Zn* | -0.01 | 0.9 | -0.07 | 0.83 | -0.1 | 0.91 | 0.01 | 0.41 |
| *Ca* | 0.04 | 0.25 | 0.03 | 0.33 | 0.06 | 0.14 | 0.07 | 0.16 |
| *Fe* | 0.01 | 0.52 | -0.03 | 0.65 | -0.1 | 0.83 | 0.01 | 0.35 |
| *Al* | **0.33** | **0.006** | **0.22** | **0.014** | **0.16** | **0.03** | **0.35** | **0.01** |
| *K* | -0.03 | 0.61 | -0.05 | 0.78 | 0.04 | 0.23 | -0.03 | 0.65 |
| *pH* | **0.12** | **0.02** | 0.07 | 0.11 | 0.07 | 0.1 | **0.14** | **0.02** |
| *Mg* | 0.01 | 0.47 | 0.03 | 0.31 | 0.05 | 0.21 | 0.02 | 0.38 |
| *Total N%* | **0.26** | **0.005** | 0.07 | 0.18 | 0.08 | 0.13 | **0.26** | **0.008** |
| *Total C%* | **0.19** | **0.017** | 0.03 | 0.3 | 0.04 | 0.31 | **0.25** | **0.005** |

## DISCUSSION

In this study, we explored the influence of multiple factors on the composition of soil fungal communities associated with tree saplings in a small woodland. It is widely accepted that many abiotic and biotic factors can influence the composition and community assembly of soil fungal taxa in temperate forests, but these can differ depending on study location and scale (Talbot et al. 2014; Tedersoo et al. 2014; Lladó et al. 2018). Our woodland experimental assay differed in tree diversity, tree compositions, and soil characteristics across space, but we found that these factors explained only a small proportion of the total variation in fungal community composition within this small woodland. Further, in the absence of broad community compositional differences, we could also identify no significant difference in extracellular enzymatic activity among treatments. However, this local woodland-based bioassay did allow for the reduction in potential confounding effects of regional and continental environmental changes, while focusing on the relative importance of tree species identity and spatial heterogeneity of soil characteristics on soil fungal community composition.

Consistent with our first hypothesis, soil fungal community composition was not significantly different between the two planted sapling species, *Quercus rubra* and *Pinus resinosa*, or their paired composition. Although there is evidence that tree seeds have their own microbiome, these results suggest that tree sapling’s fungal communities in this local woodland are obtained from a regional species pool that has already been influenced by nearby mature tree species and historic soil factors (Cline et al. 2005; Shade et al. 2017; Lladó et al. 2018). Therefore, the location where a tree seed germinates likely determines which fungal taxa associate with the sapling as it matures (Cline et al. 2005; Dickie and Reich 2005). In agreement with our second hypothesis, the data suggest that the mature trees of *Q. rubra* and *F. grandifolia* distributed in this woodland did have distinct fungal community compositions from one another. However, mature tree species identity was a weak predictor of community dissimilarity. This pattern is not unexpected given that these two tree species are in the same family (Fagaceae), and there is evidence which suggests a positive relationship between tree phylogenetic relatedness and soil fungal community similarity (Ishida et al. 2007; Lang et al. 2011; Urbanová et al. 2015). The lack of community dissimilarity between our planted sapling treatments at the phylum level is also not unexpected given that the main phyla found in the dataset are very common in temperate forests in the northeastern US and associate with many tree hosts (Tedersoo et al. 2014; Barnes et al. 2021). Compositional homogeneity in soil fungi across this woodland might itself facilitate seedling establishment that are closely related to the tree diversity aboveground creating potential positive feedback (Bennett et al. 2017).

While tree species identity was not a major factor in influencing fungal community composition, different areas of the woodland did have distinct fungal communities possibly due to variations in soil characteristics including soil moisture, total C, total N, and concentrations of Al. These influential characteristics are important to soil functionality and are linked to soil fungal distributions due to their influence on fungal physiology (Tedersoo et al. 2014; Baldrian 2017; Lladó et al. 2018). As an example, taxa within two fungal families, Mycosphaerellaceae and Auriculariaceae, were only highly relatively abundant in the lowland area of this woodland (*Block 4*) which showed variation in these characteristics compared to the other woodland areas. These fungal families have been associated with wetland soils in previous studies suggesting periodic flooding of this lowland area, along with concomitant changes to soil characteristics, might be selecting for these fungal groups (Shuhada et al. 2017). Further, RDA ordination suggests pH, Al, moisture, and Ca may be influential in structuring these communities, which may be linked to the weathering or pH buffering occurring within these soils, especially in the lowland area of the woodland (Finlay et al. 2009; Landeweert et al. 2001; Clarholm and Skyllberg 2013). Of note, these influential soil characteristics explained only a small percentage of the total community variation suggesting other abiotic factors, such as leaf litter quality, or the interaction between characteristics, are also important in influencing these communities.

After considered the evaluated deterministic factors, much of the variation in community composition was left unexplained. Therefore, one potential explanation for this unexplained variation might stochastic processes such as dispersal and ecological drift which may be important in structuring these local soil fungal communities (Dumbrell et al. 2010; Peay et al. 2012; Peay and Bruns 2014, Bahram et al. 2016; Gao et al. 2020). For instance, one possibility that might explain the relative homogenization of soil fungal community composition in this woodland might be high local fungal spore dispersal linked with habitat fragmentation limiting regional spore dispersal. The woodland at the Calder Center is categorized as a suburban forest based on local population density and percent developed landcover, and therefore fragmented by surrounding housing, building construction, and road traffic, which may be acting on these fungal communities. While there were no major patterns at the phyla level, at lower taxonomic levels (e.g., genus, ASV), ecological drift, dispersal limitation, and biotic interactions over time may become more important in driving community dissimilarity (Peay and Bruns 2014; Clemmensen 2015). Further, tree saplings can become less contingent on nearby mature trees over time, which may also increase community heterogeneity (Cline et al. 2005; Dickie and Reich 2005). Therefore, resampling these trees in subsequent years could provide a deeper view into the relationship between these tree species, soil characteristics, and the soil fungal community.

In forests, there can be a structure-function relationship between soil fungal community composition and enzymatic activity (Strickland et al. 2009; Burns et al. 2013; Kyaschenko et al. 2017). However, in contradiction of our hypothesis, the potential activities of the enzymes evaluated did not significantly differ among the treatments, mature tree species identity, or woodland location. These results might suggest that these communities have high functional redundancy regardless of any differences in community composition and a decoupling between local nutrient cycling, soil characteristics, and soil fungal community composition (Strickland et al. 2009; Brockett et al. 2012; Burns et al. 2013; Kivlin and Treseder 2014; Talbot et al. 2014). Functional redundancy in soil fungal communities can arise due to broad convergences of nutrient acquisition approaches among separate fungal groups, especially among the major phyla found in the data (Talbot et al. 2014). Although not significantly different, there was higher activity on average for all evaluated enzymes in treatments around mature *Q. rubra*. This could be related to *Q. rubra* root architecture or root exudates as well as its associated soil fungal community (Strickland et al. 2009; Burns et al. 2013; Kivlin and Treseder 2014). It is important to note that only six common enzymes were evaluated in this bioassay, which may miss other enzymatic types or their interaction effects. Also, these results focus on fungi, the main decomposers in soil, yet saprobic bacteria can degrade organic matter, which may be responsible for some of the activity found in these soils (Schneider et al. 2012; Talbot et al. 2014). Because soil enzyme dynamics are complex, caution should be applied in broader interpretation of this data, but these results do add to evidence of the factors that may or may not be influencing functional enzymatic activity in temperate forests.

Overall, this study demonstrated the roles of tree species identity and soil characteristics on the soil fungal communities of tree saplings in a small woodland. Despite the well-established relationship between fungal community composition and tree identity, the results suggest that in this woodland, these communities have assembled largely independent of the aboveground tree distribution. This has led these communities to be mainly homogeneous at broader taxonomic levels. The fungal communities of the planted saplings were different depending on the nearby mature tree species identity, but location within this woodland explained more of the variation in community composition. Additional hypotheses are needed to identify alternative factors impacting these communities as most of the variation in composition was left unexplained. Finally, the data also suggest a dissociation in the structure-function relationship between changes in fungal community composition and enzymatic activities. Linking the abiotic and biotic ecological drivers of soil fungal community composition is vital to understanding and predicting how both plants and fungal communities might be impacted by future global change.

## DECLARATIONS

## Acknowledgments

We thank the Louis Calder Center for the use of their facilities including the Calder Forest and the greenhouse. We also thank Drs. Steven Franks, John Wehr, and Patricio Meneses for their feedback on earlier drafts of this manuscript.

